# Poorer auditory sensitivity is related to stronger visual enhancement of the human auditory mismatch negativity (MMNm)

**DOI:** 10.1101/604165

**Authors:** Cecilie Møller, Andreas Højlund, Klaus B. Bærentsen, Niels Chr. Hansen, Joshua C. Skewes, Peter Vuust

## Abstract

Multisensory processing facilitates perception of our everyday environment and becomes particularly important when sensory information is degraded or close to the discrimination threshold. Here, we used magnetoencephalography and an audiovisual oddball paradigm to assess the complementary role of visual information in subtle pitch discrimination at the neural level of participants with varying levels of pitch discrimination abilities, i.e., musicians and nonmusicians. The amplitude of the auditory mismatch negativity (MMNm) served as an index of sensitivity. The gain in amplitude resulting from compatible audiovisual information was larger in participants whose MMNm amplitude was smaller in the condition deviating only in the auditory dimension, in accordance with the multisensory *principle of inverse effectiveness*. These findings show that discrimination of even a sensory-specific feature as pitch is facilitated by multisensory information at a pre-attentive level, and they highlight the importance of considering inter-individual differences in uni-sensory abilities when assessing multisensory processing.

## Poorer auditory sensitivity is related to stronger visual enhancement of the auditory mismatch negativity (MMNm)

Because the world is multimodal, everyday perception relies on the complementary roles of simultaneously active sensory systems. Contemporary brain science focuses increasingly on the principles underlying cortical connectivity (Friston, 2005), and the notion that multisensory processing in the brain only occurs after extensive processing in strictly sensory-specific cortical areas (Felleman & Van Essen, 1991) has been challenged in the past decades (for a classic review, see Ghazanfar & Schroeder, 2006), where an increasing number of studies have shown crossmodal influences on processing in primary sensory areas such as the auditory cortex (Atilgan et al., 2018; Brosch, Selezneva, & Scheich, 2005; Jäncke, Rogenmoser, Meyer, & Elmer, 2012; van Wassenhove & Grzeczkowski, 2015).

The *mismatch negativity* (MMN) and its magnetic counterpart MMNm (also termed *mismatch field* or *mismatch response*), is used to probe pre-attentive levels of unisensory auditory processing (Näätänen, Paavilainen, Rinne, & Alho, 2007). It is a robust measure derived from the event-related potential (ERP) / event-related field (ERF), which can be elicited in the auditory cortex in response to deviants in a repetitive train of auditory tones, and it is regarded as an index of various auditory functions, including neural sound discrimination and sensitivity (Näätänen, 2008). The amplitude of the MMN is positively correlated with the magnitude of the deviant (Jaramillo, Paavilainen, & Näätänen, 2000; Sams, Paavilainen, Alho, & Näätänen, 1985) as well as with behavioral performance (Amenedo & Escera, 2000; Jaramillo et al., 2000; Novitski, Tervaniemi, Huotilainen, & Näätänen, 2004; Tervaniemi, Just, Koelsch, Widmann, & Schroger, 2005), which makes the MMN paradigm a useful method to assess perceptually relevant neural sensitivity to subtle changes in an auditory stimulus material.

Nonetheless, MMN/MMNm elicitation is not restricted to unisensory auditory processing as strong audiovisual illusions may also elicit an MMN/MMNm in the auditory cortex. A long tradition of scientific studies has used crossmodal perceptual illusions to elucidate the importance and the consequences of the spontaneous cooperation between the sensory modalities at behavioral (McGurk & MacDonald, 1976; Sekuler, 1997; Shams, Ma, & Beierholm, 2005) and neural levels (Bonath et al., 2007; Colin, Radeau, Soquet, Dachy, & Deltenre, 2002; Colin, Radeau, Soquet, Demolin, et al., 2002; Möttönen, Krause, Tiippana, & Sams, 2002; Saint-Amour, De Sanctis, Molholm, Ritter, & Foxe, 2007; Sams et al., 1991; for review, see Alais, Newell, & Mamassian, 2010).

For instance, in the McGurk effect (McGurk & MacDonald, 1976), a mismatch between visual (ga-ga) and auditory (ba-ba) speech phonemes most often results in a perceptual experience of a syllable (da-da) which is not a veridical representation of either of the sensory constituents. In a pioneering magnetoencephalography (MEG) study, Sams et al. (1991) used the McGurk effect to show visual modulation of mismatch responses elicited in the auditory cortex in response to deviant perception of identical sounds. Kislyuk, Möttönen, and Sams (2008) complemented the study by Sams et al (1991) by showing that the MMN elicited in response to deviant speech sounds (‘ba’ in a train consisting of ‘va’) may be suppressed by simultaneous presentation of incongruent visual (‘va’) information. Notably, in both of these experiments, the auditory stimuli were kept constant but generated different mismatch responses across conditions. This highlights the link between MMN elicitation and perceptual experience over and above physical properties of the auditory stimuli.

A number of studies have assessed the relevance of congruent visual information to non-illusory speech processing in the auditory cortex, (Calvert et al., 1999; Möttönen, Schürmann, & Sams, 2004; Okada, Venezia, Matchin, Saberi, & Hickok, 2013; Pekkola et al., 2005; Smith et al., 2013). Comparatively less is known about crossmodal influences on cortical processing of more basic auditory features such as pitch. This is noteworthy considering the longstanding and recently intensified scientific interest in crossmodal correspondences between auditory pitch and visual features such as brightness (Klapetek, Ngo, & Spence, 2012; Marks, 1987), lightness (Melara, 1989; Melara & O’Brien, 1987), angularity (Marks, 1987; Walker, 2012), size (Evans & Treisman, 2010; Gallace & Spence, 2006), and vertical position (Bernstein & Edelstein, 1971; Deroy & Spence, 2016; Melara & O’Brien, 1987; Møller et al., 2018; Orchard-Mills, Van der Burg, & Alais, 2015; Parise, Knorre, & Ernst, 2014; Pratt, 1930; see also Spence, 2011). In a series of studies, Besle and colleagues used electroencephalography (EEG) (Besle et al., 2007; Besle, Fort, & Giard, 2005; Besle, Hussain, Giard, & Bertrand, 2013) and MEG (Besle et al., 2007) to study visual influences on the MMN(m) elicited in response to salient pitch changes. These studies indicated that visual stimuli have access to the processes in the auditory cortex that underlie the generation of MMN(m) responses elicited to non-speech and non-illusory stimuli, specifically *salient* pitch changes. If that is the case, then visually induced behavioral facilitation of *subtle* pitch discrimination should also be evident at the pre-attentive stage of processing probed by the MMNm.

We recently conducted an audiovisual behavioral experiment that capitalized on the inter-individual variability that characterizes pitch discrimination. We showed that the size of visually induced gain in subtle pitch discrimination at the behavioral level is inversely related to participants’ musical aptitude and auditory pitch discrimination skills as measured by unisensory tools (Møller et al., 2018). This finding is consistent with an *inter-individual* interpretation of the multisensory *principle of inverse effectiveness* (PoIE) according to which, multisensory integration is maximal when responses to the unisensory dimensions of the multisensory event are weak (Stein & Meredith, 1993). The functional relevance of this principle is obvious: when a unisensory dimension of a multisensory event is hardly detectable, verification through other sensory modalities is essential. Notably, previous work in support of PoIE at behavioral and neural levels assessed *intra-individual* effects, i.e., by comparing group-averaged responses to varying levels of stimulus salience (Senkowski, Saint-Amour, Hofle, & Foxe, 2011). In our behavioral study, we also found a significant congruence effect, i.e., participants performed better when crossmodally matching than mismatching visual cues were presented simultaneously with target pitch deviants. The question remains whether a similar *inter-individual* interpretation of the PoIE and the congruence effect are evident at the pre-attentive stage of pitch processing probed by the MMNm or whether later levels of processing (e.g., semantic or decisional processes) are exclusively responsible for the pattern of behavioral benefits observed.

Consequently, in the present MEG study we asked whether the visual dimension of audiovisual stimuli could modulate the MMNm elicited in the auditory cortex in response to identical subtle pitch changes, and if so whether the size of the visual modulation depends on the degree of unisensory expertise at the level of the individual. This focus necessitated recruitment of participants spanning a wide range of pitch discrimination abilities. Motivated by previous studies that have shown enhanced MMN amplitudes in musicians compared to nonmusicians (Tervaniemi et al., 2005; Vuust, Brattico, Seppanen, Naatanen, & Tervaniemi, 2012), a subgroup of musicians were added to the participant pool.

We modified the audiovisual oddball paradigm used in the previous behavioral study, allowing us to quantify visually induced gain at the neural level by comparing the amplitude of MMNm responses elicited to deviants differing from the standards in the auditory dimension (pitch), the visual dimension (vertical position), or both dimensions. Notably, when both dimensions varied, they did so in the same or opposite directions of each other, according to the crossmodal correspondence between a high pitch and a high vertical position (Spence, 2011), as this allowed us to assess the effect of crossmodal congruence on pre-attentive pitch processing. To testify to the applicability in the present study of the MMNm amplitude as an index of neural pitch discrimination sensitivity, we also compared individual measures of MMNm amplitude in all conditions against behavioral measures of pitch discrimination thresholds and of musical aptitude.

These steps were taken towards the main goal of the study, which was to assess inter-individual differences in audiovisual pitch processing. We tested the specific hypothesis that larger visually induced MMNm enhancements would be present in participants with smaller MMNm responses elicited to auditory-only pitch deviants, (i.e., deviants presented without any complementary visual information). This would be consistent with an inter-individual interpretation of the PoIE and with our previously reported behavioral results (Møller et al., 2018), and it would highlight the need for taking individual differences in unisensory neural sensitivity into accout when assesing multisensory processing.

## Results

### Group-level results

Differences in MMNm amplitudes in response to the four conditions *same direction* deviants (AVsame), *opposite direction* deviants (AVopp), *auditory-only* deviants (Adev), and *visual-only* deviants (Vdev) were assessed separately in the two hemispheres using Friedman ANOVAs. Exact statistics are reported. There was a statistically significant main effect of condition in the left, ***χ***^***2***^(3) = 21.507, *p* < .001, as well as in the right hemisphere, ***χ***^***2***^(3) = 20.520, *p* < .001. Boxplots showing medians and the interquartile ranges of responses in the different conditions are reported in Figure 1, along with plots showing the average ERF waveforms elicited by the standard and the four deviants. The distributions of individual MMNm responses to all conditions are shown in supplementary materials, Figure S1. Group-averaged gradiometer and magnetometer topographies are also reported in supplementary materials, Figure S2, S3, S4, and S5).

**Figure 1.**
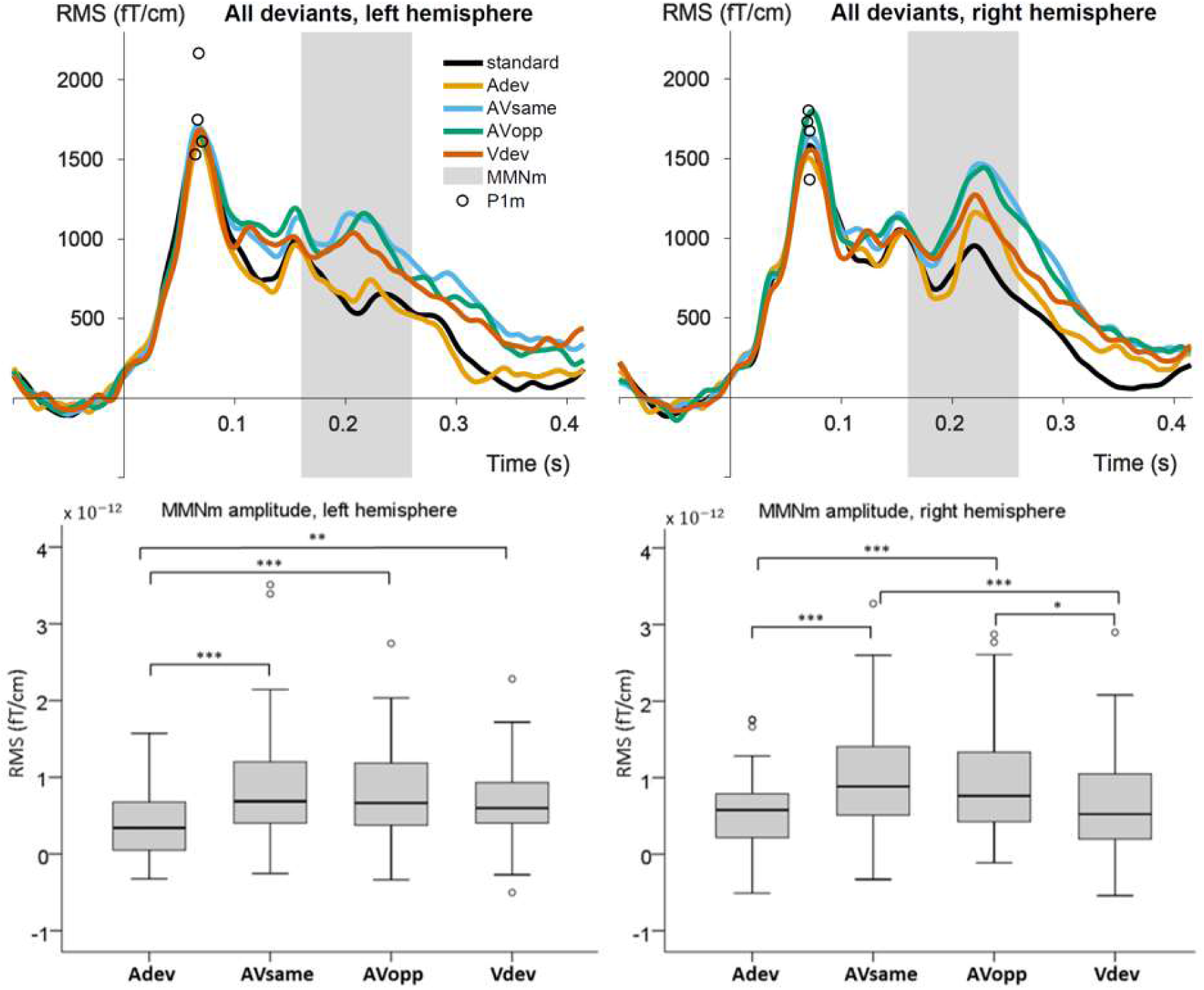
Top: Group-averaged ERF waveforms elicited by the standard and the four deviants. Note, that statistics were performed on the amplitudes of MMNm responses identified in each individual participant (see individual responses to each deviant in supplementary materials, Figure S1). Circles show amplitudes and latencies of the P1m peaks in the four channels used for analysis in each hemisphere. Bottom: Boxplots show medians and the interquartile ranges of responses (individually identified MMNm responses) in the different conditions. * *p <* .008, ** *p <* .00167, *** *p <* .000167

Statistical significance of the subsequent six pairwise comparisons was assessed with Wilcoxon Signed-Ranks tests at a Bonferroni-corrected significance level of α = .008. *Z* statistics, asymptotic *p*-values, and effect sizes are reported in Table 1. Importantly, significantly larger MMNm amplitudes were observed in both hemispheres in response to AVsame compared to Adev, consistent with the hypothesis. However, the present analysis did not show evidence that congruity between the auditory and visual components of the stimuli had an effect on the size of the MMNm enhancement, as there was no significant difference between responses to AVsame and AVopp stimuli. Notably, despite no pitch change, an MMNm-like component was observed above the auditory cortex in response to Vdev stimuli. In the right hemisphere, the response was significantly smaller than that elicited by AVsame and AVopp, but the size of it was not significantly different from that elicited by Adev stimuli. In the left hemisphere, the response was significantly larger than the Adev MMNm, but it was not significantly different from that elicited by the AVsame and AVopp stimuli.

**Table 1.** Pairwise comparisons

| Comparison | *Z* |  | *p* |  | *r* |  |
| --- | --- | --- | --- | --- | --- | --- |
|  | Left | Right | Left | Right | Left | Right |
| AVsame vs. Adev | -4.442 | -3.798 | < .001*** | < .001*** | -.47 | -.40 |
| AVsame vs. AVopp | -0.141 | -0.751 | .888 | .453 | -.001 | -.08 |
| Adev vs. Vdev | -3.200 | -0.528 | .001 ** | .600 | -.34 | -.06 |
| AVopp vs. Adev | -4.295 | -3.685 | < .001 *** | < .001*** | -.45 | -.39 |
| AVsame vs. Vdev | -1.180 | -4.058 | .238 | < .001*** | .12 | -.43 |
| AVopp vs. Vdev | -1.022 | -2.749 | .307 | .006* | -.11 | -.29 |

*Notes.* Wilcoxon Signed-Ranks tests assessed the pairwise comparisons between conditions.
Adev = auditory-only deviant, AVsame = audiovisual, same direction, AVopp = audiovisual, opposite direction, Vdev = visual-only deviant. Asymptotic *p*-values were assessed at Bonferroni-corrected significance levels. \* *p* < .008, \*\* *p* < .00167, \*\*\* *p* < .000167.

### Visually induced enhancements and unisensory skills

Visually induced MMNm-enhancement was defined as the difference in MMNm amplitude elicited by AVsame compared to Adev stimuli, henceforth denoted ΔMMNm. Behaviorally measured pitch discrimination thresholds did not predict ΔMMNm, neither in the left hemisphere, *r* = -.237, 95% CI [-.453, .067], nor in the right hemisphere, *r* = -.047, 95%, 95% CI [-.258, .364], as assessed with Pearson correlation analyses with bootstrapped bias-corrected and accelerated (BCa) 95% confidence intervals (5000 samples). However, the amplitude of MMNm responses elicited to Adev stimuli were statistically significantly inversely related to ΔMMNm, i.e., larger Adev amplitude was associated with smaller ΔMMNm, *r* = -.397, 95% CI [-.595, -.194], in the right hemisphere. See scatterplots in Figure 2. Although no such statistically significantly association was found in the left hemisphere, *r* = -.183, 95% CI [-.399, .051], the correlation was significant when responses of the left and right hemisphere were averaged, (see supplementary materials, Figure S6).

**Figure 2.**
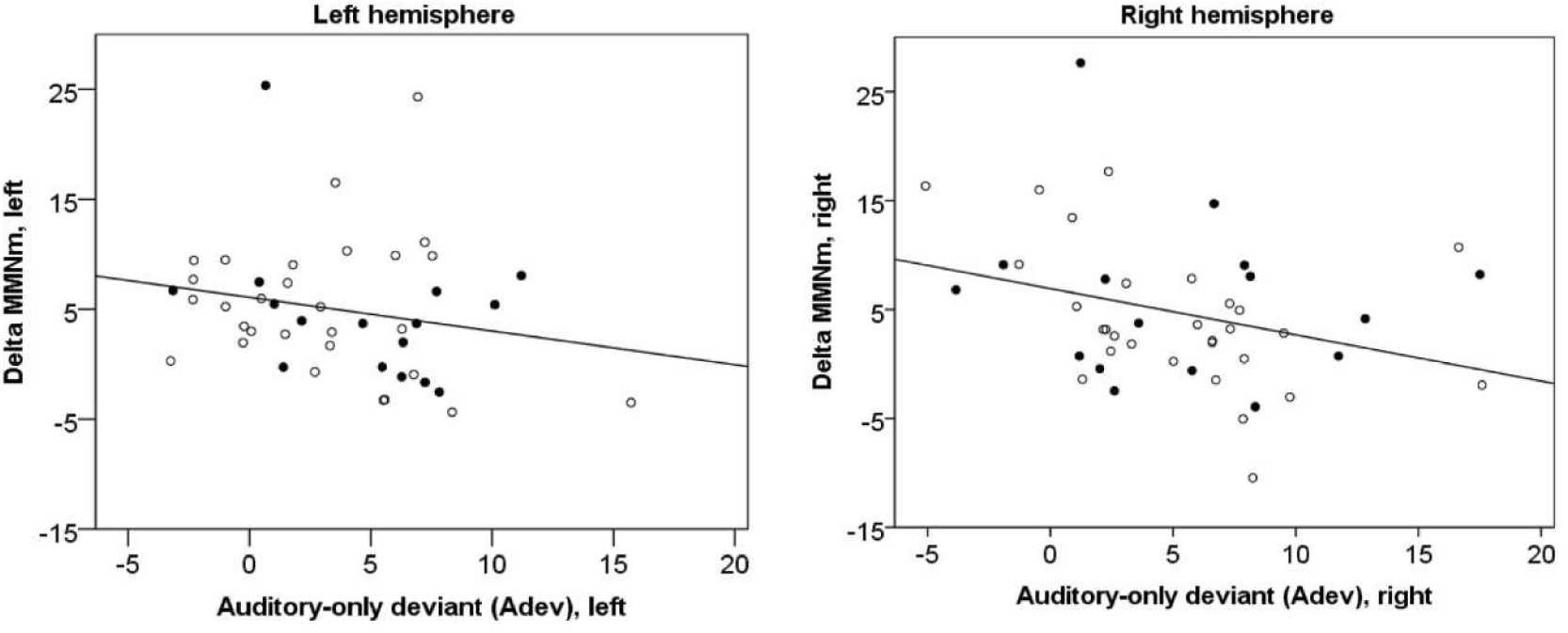
Plots show the visually induced enhancement in MMNm amplitude, ΔMMNm, as a function of the MMNm amplitude elicited in response to auditory-only deviants, Adev, i.e., without complementary visual cues. The correlation is significant in the right hemisphere as assessed with bootstrapped bias-corrected and accelerated (BCa) 95% confidence intervals (5000 samples). Solid circles = musicians, open circles = nonmusicians

### Behavioral correlates to MMNm amplitude

Bootstrapped Pearson’s correlation analyses were run to examine the relationship between MMNm amplitude and behavioral scores obtained in the Musical Ear Test (MET) (Wallentin, Nielsen, Friis-Olivarius, Vuust, & Vuust, 2010) and the pitch discrimination threshold (PDT) estimation. Statistical significance was assessed at α = .05. PDT data points were nonnormally distributed and hence log-transformed using the natural logarithm. Left-hemisphere MMNm amplitudes in response to all four experimental conditions were significantly correlated with PDT and with MET(melodic) scores, i.e., larger MMNm amplitudes were associated with lower (i.e., better) pitch discrimination thresholds and with better performance on the MET, testifying to the applicability in the present study of the MMNm amplitude as an index of neural pitch discrimination sensitivity. No significant associations were found between behavioral measures and MMNm amplitudes elicited in the right hemisphere. See Figure 3 for scatterplots and Table 2 for correlation coefficients and bootstrapped bias-corrected and accelerated (BCa) 95% confidence intervals. A Pearson product-moment correlation analysis showed that MET and PDT scores were also significantly correlated (*r* = -.528, *p* <.001), reflecting the association between musical skills and auditory sensitivity.

**Table 2.**
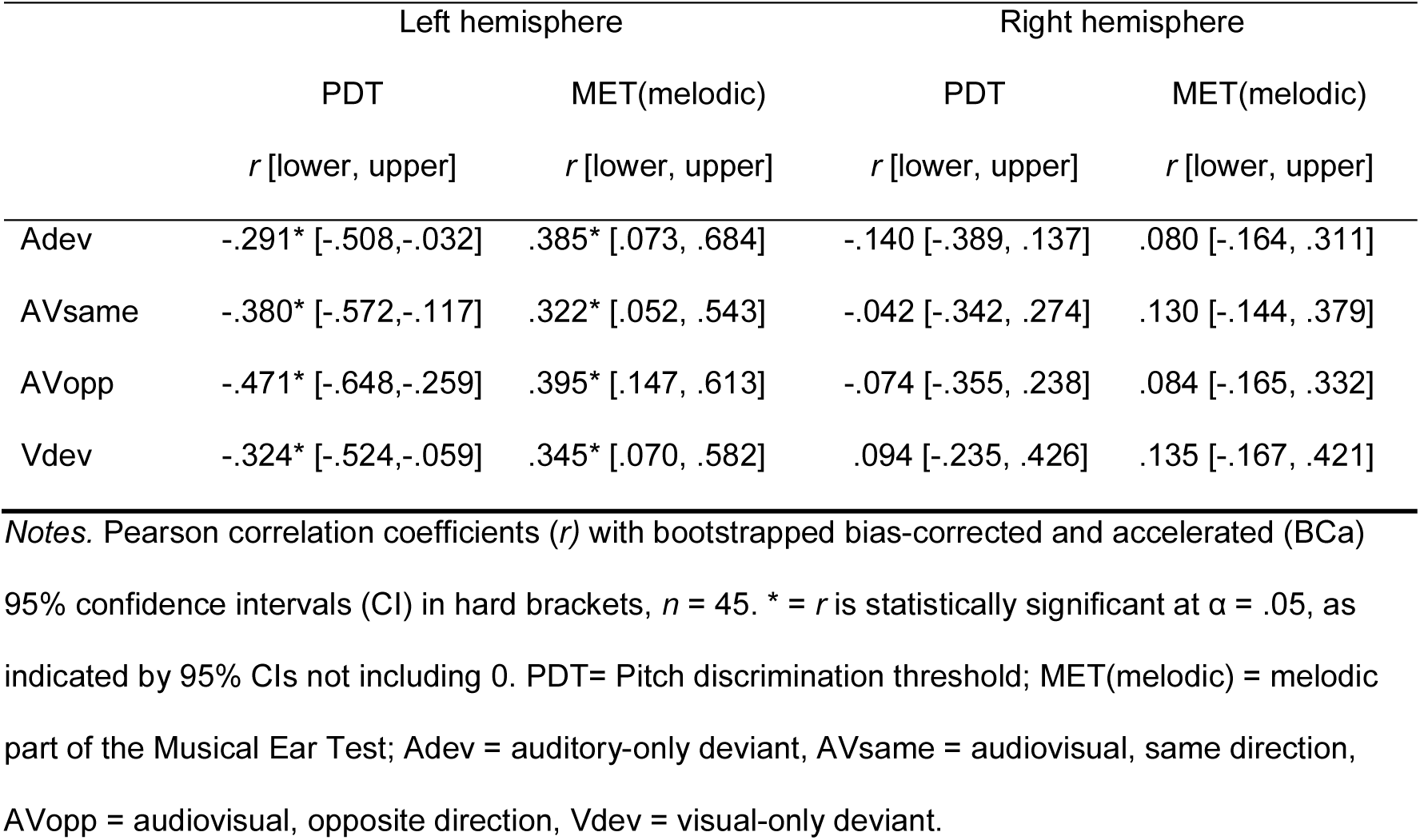
Correlations between behavioral measures and MMNm amplitudes in the four conditions

**Figure 3.**
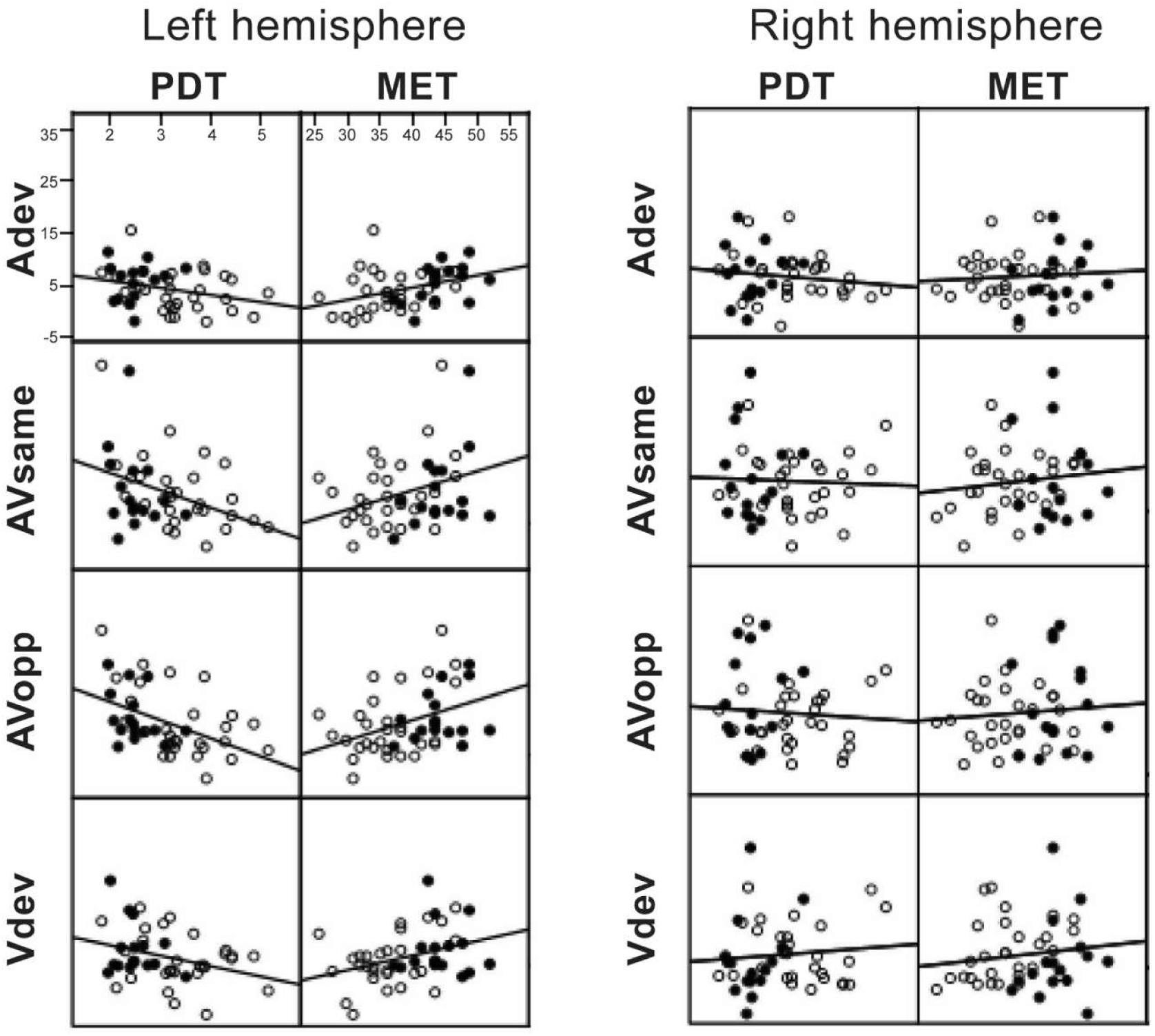
Scatterplots show MMNm amplitudes (y-axes) plotted against behaviorally measured log-transformed pitch discrimination thresholds (PDT) and behavioral scores on the melodic part of the Musical Ear Test (MET). For all conditions, the correlation is significant in the left hemisphere as assessed with bootstrapped bias-corrected and accelerated (BCa) 95% confidence intervals. PDT= Pitch discrimination threshold; MET(melodic) = melodic part of the Musical Ear Test; Adev = auditory-only deviant, AVsame = audiovisual, same direction, AVopp = audiovisual, opposite direction, Vdev = visual-only deviant. Solid circles = musicians, open circles = nonmusicians

## Discussion

This study was designed to assess inter-individual differences in visually induced enhancements of subtle pitch discrimination at the pre-attentive stage of perceptual processing. Across all participants (musicians and nonmusicians), the amplitude of the MMNm recorded using MEG was significantly enhanced in conditions containing audiovisual deviants (AVsame and AVopp) compared to a condition in which only the auditory dimension of the audiovisual stimuli deviated (Adev). In the right hemisphere, the size of this enhancement was significantly correlated with the amplitude measured in the Adev condition, indicating that the visual dimension of audiovisual events has a larger impact on pre-attentive pitch discrimination in the auditory cortices of people with lower auditory-only pitch discrimination sensitivity. This result is consistent with an inter-individual interpretation of the *principle of inverse effectiveness* (PoIE) and suggests not only that individual differences in multisensory processing exist, but also that they may be explained in part by individual differences in unisensory sensitivity.

Our study contributes to the literature showing that processes occurring early in sensory-specific cortices can be modulated by crossmodal information (see also Bulkin & Groh, 2006; Driver & Noesselt, 2008; Ghazanfar & Schroeder, 2006). We focused specifically on visually induced enhancements of pitch discrimination in the auditory cortex. It is important to note that perceptual enhancement is but one of several possible outcomes of multisensory integration (Driver & Noesselt, 2008) and also that the MMN(m) has been interpreted specifically as indexing the ability of the brain to detect incongruent relations between auditory and visual stimuli, i.e., as an incongruency response (Paraskevopoulos, Kuchenbuch, Herholz, & Pantev, 2012b). Our interpretation of the MMNm as an index of perceptual sensitivity rather than an incongruency response rests on previous studies showing that the MMN is related to behavioral performance in tasks that employ similar sounds (Jaramillo et al., 2000; Novitski et al., 2004) and in groups of participants with different levels of auditory abilities, i.e., musicians and nonmusicians (Tervaniemi et al., 2005). This is also confirmed in our data where the correlations between MMNm amplitudes and behavioral measures of pitch discrimination abilities and musical aptitude testify to the applicability of the MMNm amplitude as an index of pre-attentive neural pitch discrimination.

### Group-level results

Visually induced MMNm enhancement was evident in all conditions of the experimental paradigm in which the visual dimension of the audiovisual event deviated (AVsame, AVopp, Vdev) compared to the condition in which it did not (Adev). Our primary interest was the assessment of the difference in MMNm amplitude elicited by AVsame compared to Adev stimuli, i.e., ΔMMNm. This was the most important outcome variable as the key objective of the study was to assess whether this reliance followed the PoIE at the inter-individual scale, i.e., varied with varying levels of auditory-only pitch discrimination abilities. At the group level, this difference was highly significant in both hemispheres, consistent with the notion that visual information not only induces a qualitatively different perceptual experience and hence elicits an MMNm (Möttönen et al., 2002; Saint-Amour et al., 2007; Sams et al., 1991), but that it also modulates the size of the MMNm (Besle et al., 2013) elicited to identical auditory deviants. A discussion of the individual-level results will follow after further considerations of the group-level results.

The comparison between AVsame and AVopp constitutes what we have previously referred to as the crossmodal congruence effect (Møller et al., 2018). Note, that we used the feature-based crossmodal correspondence between auditory pitch and visually perceived vertical position because we sought to maximize the likelihood that the visual stimuli would be beneficial. In this perspective, the disentanglement of two kinds of facilitatory effects (i.e., ΔMMNm and congruence effect) was not as much an aim as a by-product of the paradigm, caused by the polarity inherent in the audiovisual stimuli, where the polar dimensions of auditory pitch and visually perceived vertical position map onto each other. From this also follows that we refer to crossmodal congruence in relative terms, in contrast to studies that conceptualize audiovisual congruence in absolute terms, i.e., violations of absolute distances in the same direction are defined as audiovisual incongruence (Nichols & Grahn, 2016) or no change in the visual dimension (comparable to our Adev) is considered an incongruent trial(Paraskevopoulos, Kuchenbuch, Herholz, & Pantev, 2012a). To avoid confusions with the inconsistent use of the terms evident in the literature, we refrain from using the labels congruent and incongruent to denote trials, but use congruence effect merely to denote the comparison between AVsame and AVopp. We did not find a significant congruence effect in the present study, despite the fact that we did so in our previously reported behavioral study using similar stimuli. The most obvious candidate explanation for this is that the feature-based crossmodal correspondence used does not influence pre-attentive levels of perceptual processing probed by the MMNm. This interpretation of the result taps into the debate about the level at which these crossmodal interactions take place (Deroy & Spence, 2016; Parise & Spence, 2009) and could be taken as an argument that this specific correspondence only influences participants’ responses at semantic and/or decisional levels of processing. Yet, first of all, one should be careful of drawing conclusions based on null-results, and secondly, there is still the possibility that less salient visual stimuli than the ones used in the present paradigm would lead to significant congruence effects, as bimodal response enhancement is most pronounced when both stimulus dimensions in isolation lead to weak responses, according to the PoIE (Stein & Meredith, 1993). As is evident in the topographies of the conditions involving changes in the visual stimuli, this is clearly not the case here (see supplementary materials, Figure S2, S3, S4, S5).

Although the visual-only deviant (Vdev) was not an explicit focus of our study, as explained in the methods section, we briefly provide two possible yet still speculative interpretations of the finding that an MMNm comparable in size to the Adev (right hemisphere) and AVsame and AV opp (left hemisphere) was elicited to Vdev, despite no change in the auditory dimension of the stimulus. The first is based on a simple idea of classical conditioning of the auditory cortex (Pavlov, 1927). Because all conditions were included in the same paradigm, it is possible that the MMNm response to Vdev was a conditioned reflex, i.e., the result of a conditioning process in which the auditory and visual dimensions of the audiovisual stimulus have been repeatedly coupled. Crossmodal conditioning and anticipatory excitation of the cortex was demonstrated already in the early 1940’s (Anokhin, 1974) and more recent findings also support the notion that the perceptual system easily adapts its predictions even to unnatural stimulus events through the course of an experiment (van Laarhoven, Stekelenburg, & Vroomen, 2017). Indeed, in the visual modality it has been shown that after a visuohaptic training session, haptic shape processing activates visual cortex in the absence of visual input (Lee Masson, Bulthé, Op de Beeck, & Wallraven, 2016).

An alternative, yet most likely related account, may be rooted in the fundamental object-directed nature of perception, and the fact that spatiotemporal coherence is key to multisensory object formation processes (Atilgan et al., 2018; Lewkowicz & Ghazanfar, 2009; Parise, Spence, & Ernst, 2012). Although we presented participants with simple artificial auditory and visual stimuli, their consistent temporal synchronicity may have made them so convincing audiovisual objects, that an MMNm was elicited whether the auditory, the visual, or both dimensions of the audiovisual stimulus deviated. MMNm elicitation is not restricted to conditions in which the physical properties of the sounds deviate, but may also be elicited to violations of abstract feature patterns (Näätänen et al., 2007). Assuming the perceptual system recognizes the stimuli as integrated audiovisual percepts, it is not surprising that a violation in any of its dimensions give rise to an abstract-feature MMNm. Validation of such an account, however, would require demonstrations that similar processes occur in the visual cortices, which is beyond the scope of the present study.

### Individual-level results

This is the first study to quantify the size of visually induced MMNm enhancement across participants with varying levels of unisensory auditory abilities in order to show that the MMNm amplitude enhancement is larger in individuals with smaller amplitudes elicited to auditory-only deviants. This result matches our previously reported behavioral findings of larger gains in pitch discrimination with poorer pitch discrimination thresholds (Møller et al., 2018), and is consistent with an inter-individual interpretation of the PoIE.

Similar patterns of observation have been found at the intra-individual level in studies investigating vision as the target modality. Noesselt and colleagues (2010) employed group-level analyses of fMRI data to show sound-induced sensitivity enhancements to low-intensity but not high-intensity visual stimuli. Stevenson and colleagues (2012) recorded visual ERPs across salience levels of audiovisual speech and found greater multisensory gains as salience was reduced. The role of uni-sensory expertise was assessed by Caclin and colleagues (2011) who investigated sound-induced enhancements of visual detection thresholds and found that behavioral enhancements were only evident in a group of participants with poor performance in a visual-only condition. Similarly, in the auditory domain, visual facilitation of behavioral pitch discrimination of quite salient pitch changes (1/4 tone, i.e., 50 cents) has been found only in a group of participants with poor auditory-only abilities (Albouy et al., 2015), while conversely, only the control group exhibited visual facilitation of the subtle (1/16 tone, i.e., 12.5 cents) pitch changes. Senkowski and colleagues (2011) studied effects of stimulus intensities on both auditory and visual modalities and found evidence that early multisensory interactions in the human brain follow the PoIE. Our results extend these group-level findings to show that the PoIE is also applicable to pre-attentive auditory processing at the inter-individual level, highlighting the value of taking individual differences in unisensory abilities into account when assessing multisensory perceptual processing.

### Principle of inverse effectiveness

Though the PoIE was derived from studies of single multisensory neurons in the midbrain (Stein & Meredith, 1993), it not surprising to find inverse effectiveness as a general principle across a wide range of processing levels and functions (Albouy et al., 2015; Caclin et al., 2011; Møller et al., 2018; Noesselt et al., 2010; Senkowski et al., 2011; Stevenson et al., 2012). This is the case because multisensory processing does not take place in individual neurons or even cortical areas in isolation. The development of the multisensory processing capabilities of individual neurons in the superior colliculus depend on their embeddedness in a neural network consisting of lower-order subcortical areas as well as higher-order association cortices, and on the organism’s experience in the environment (Stein, Stanford, & Rowland, 2014).

Similarly, our findings of inverse effectiveness at the pre-attentive level of neural processing in the auditory cortex should not be taken to suggest, that other areas of the brain do not play a role in the kind of multisensory processing studied here. Rather, the interpretation put forth here is similar to previous accounts of multisensory influences on sensory specific cortices that highlight the fundamental object-directed nature of perception and how object formation processes in the brain are made possible through the spatio-temporal coherence of crossmodal information belonging to the same event (Atilgan et al., 2018; Lewkowicz & Ghazanfar, 2009; Parise et al., 2012). Specifically, it has been shown that phase resetting of ongoing neuronal oscillations is a key mechanisms underlying the amplification of neuronal responses. This phase resetting occurs when bimodal stimuli are temporally aligned, and ensures that sensory-specific information arrives in sensory-specific cortices during a high excitability phase resulting in perceptual amplification (Atilgan et al., 2018; see also Lakatos, Chen, O’Connell, Mills, & Schroeder, 2007; Maddox, Atilgan, Bizley, & Lee, 2015). As such, it is not the visual information itself that gives rise to excitation of neurons in the auditory cortex, but rather the excitability of the auditory neurons that is enhanced, mediated by the crossmodal synchronicity. “Top-down” influences on sensory-specific cortices are thus crucial in amplifying the salience of the sensory-specific information, in this case leading to the experience of improved hearing.

The design of this study was originally motivated by the PoIE, however, the results may also be interpreted within theoretical accounts that consider multisensory percepts as a weighted average of sensory estimates (Ernst & Banks, 2002; Schwartz, 2010). Like the PoIE, these models highlight the flexibility of our perceptual systems to adapt to variations in the sensory environment (darkness, noise). It follows, that if the reliability of the auditory dimension of the audiovisual events in our experiment is considered low in some participants and high in others, then they will give more weight to the visual and auditory dimensions, respectively. This is in line with recent evidence that musicians who specialize in the auditory domain are less susceptible to audiovisual illusions (Bidelman, 2016; Proverbio, Massetti, Rizzi, & Zani, 2016). Our finding that such effects are evident in the MMNm amplitude fits well with recent research showing that reliability-based cue weighting occurs early (< 150ms) in the audio-visual integration process (Boyle, Kayser, & Kayser, 2017), i.e., well before the latencies of the MMNm responses in the present study.

Although the correlations betwee(Alho et al., 1996; Giard & Peronnet, 1999; Tervaniemi et al., 1999)n ΔMMNm and Adev showed similar trends in the two hemispheres, it was only statistically significant in the right hemisphere. A possible explanation for this discrepancy could be rooted in the fact that larger amplitudes were seen in the right than in the left hemisphere. It may be the case that these larger amplitudes enabled detection of a significant correlation in the right hemisphere, whereas due to response homogeneity in Adev in the left hemisphere, there may not have been enough variance to detect a statistically significant correlation.

### Analysis limitations

Explanations for functional hemispheric asymmetries are highly relevant to discuss, particularly for auditory processing in the human brain. Apart from merely statistical accounts, however, there are notable reasons why further interpretations regarding lateralization can and should not be made based on the analyses reported here. Most importantly, we ran separate analyses in the two hemispheres, leaving interpretations of differences in evidence between them unsupported. Moreover, while it has been repeatedly shown that the MMN elicited to non-speech auditory stimuli is more pronounced in the right hemisphere of the general population (Alho et al., 1996; Giard & Peronnet, 1999; Tervaniemi et al., 1999), and particularly so for change detection (Levänen, Ahonen, Hari, McEvoy, & Sams, 1996), it has also been shown that expert musicians exhibit the reverse pattern, i.e., larger MMNm amplitudes in the left hemisphere (Herholz, Lappe, & Pantev, 2009; Kuchenbuch, Paraskevopoulos, Herholz, & Pantev, 2012; Vuust et al., 2005). Furthermore, Foxton and colleagues (2009) provided evidence that people with difficulty in pitch processing show greater recruitment of the left than the right hemisphere when performing active determination of pitch glide direction, a finding which may explain their poorer behavioral performance, which in turn was also evident at the pre-attentive level as assessed with the MMNm. Clearly, effects of expertise are likely to interact with any lateralization patterns evident in the present data, considering the fact that musicians and nonmusicians were not separated in the present analysis. Again, the reason for running the analyses across all participants was to ensure sufficient variance in the data to investigate individual differences, a focus that was prioritized over that of lateralization. For transparency, we have reported one group-level analysis for each hemisphere. Note, however, that when the reponses were averaged across hemispheres within participants, a procedure justified by the particular focus on individual differences, the correlation between ΔMMNm and Adev was statistically significant, consistent with the PoIE (see supplementary materials, Figure S6).

We acknowledge that a sensor-space analysis limits the conclusions we can make regarding the sources of the neural activity measured. However, there is little doubt that the MMNm is elicited in the auditory cortex, and maximally detected with gradiometers right above the local cortical source (Levänen et al., 1996; Luck, 2005; Näätänen et al., 2007), which was also confirmed in the present data at the individual as well as the group level. Furthermore, a recent study showed that MMNm detectability at the individual level is similar whether sensor-space or source-space analyses are employed. Hence, a sensor-space analysis constrained to the four gradiometer pairs of each hemisphere that picked up the largest MMNm response was considered appropriate in the present context, considering the focus on the MMNm and the fact that the MMN is very well described in the literature as a component with primary sources in the auditory cortices (Alho, 1995; Näätänen et al., 2007).

## Conclusion

Most research on perceptual processing involves stimulation of a single sense. Yet, even perceptual experiences that we consider sensory-specific, such as hearing, are influenced by information in other sensory modalities. We have shown visually induced enhancement of subtle pitch discrimination already at the pre-attentive stage of auditory processing probed by the MMNm. Importantly, participants with lower sensitivity to auditory-only deviants gain more from complementary visual information, in accordance with the multisensory *principle of inverse effectiveness.* In the quest towards understanding hearing as it unfolds outside the lab, we may gain substantially by paying attention to the possibility that hearing is related in systematic ways to other factors than sound waves.

## Materials and methods

The present magnetoencephalography (MEG) experiment forms a separate part of a larger study. In the larger study, the same participants subsequently completed a behavioral experiment (reported in Møller et al., 2018) and participated in a study employing magnetic resonance imaging (MRI) and diffusion tensor imaging (DTI) (manuscript in preparation). Behavioral measures included pitch discrimination threshold (PDT) estimations, a musical aptitude test, i.e., the Musical Ear Test (MET) (Wallentin et al., 2010), two questionnaires, i.e., the Goldsmiths Musical Sophistication Index (Gold-MSI) (Müllensiefen, Gingras, Musil, & Stewart, 2014), and the Autism Spectrum Quotient (AQ) (Baron-Cohen, Wheelwright, Skinner, Martin, & Clubley, 2001), as well as a custom-made test of pitch direction sensitivity. We report results based on the MEG data and correlate these to the behavioral measures from the MET and PDT tests. MET and PDT data were also analyzed in relation to our previously reported behavioral experiment (Møller et al., 2018), however using a smaller sample of the participants. The PDT data were also analyzed in conjunction with the DTI data (manuscript in preparation).

No explicit power analysis was used to compute appropriate sample size due to the novelty of the specific topic under study. However, sample size was established by comparisons with sample sizes used in behavioral studies of crossmodal gains and inverse effectiveness (Albouy et al., 2015; Caclin et al., 2011; Laurienti, Kraft, Maldjian, Burdette, & Wallace, 2004; Senkowski et al., 2011), where reasonable effect sizes were found with sample sizes between 11 and 34. Accordingly, we considered recruitment of ~50 participants appropriate as we expected a somewhat high exclusion rate in the subsequent behavioral study due to chance and ceiling level performance.

### Participants

MEG data were recorded from 48 participants. Data from one participant were discarded from all analyses due to stimulus presentation failure. Because of the present focus on unisensory abilities, an *a priori* decision was made to exclude data from participants from whom no pitch threshold estimation was made. This was the case for two individuals. Hence, data from 45 participants (mean age = 24.2 years, *SD* = 3.6, 23 female) were included in the analyses reported here. As described in the introduction, a sub-group of these were professional musicians. Specifically, participants included 29 nonmusicians (mean age = 24.1 years, *SD* =3.3, 16 female) and 16 musicians (mean age = 24.3 years, *SD* = 4.1, 7 female). All were right-handed, all reported normal or corrected-to-normal visual acuity and no hearing impairments.

Participants gave their written consent before participation and received a taxable compensation of DKK 400 for participating in the larger study which took place on two separate days. The study protocol was approved by The Central Denmark Regional Committee on Health Research Ethics (Project-ID: M-2014-52-14).

### Stimuli and paradigm

The auditory stimuli consisted of sinusoidal tones of 100 ms duration, including 5 ms fade in/out followed by a 316 ms inter-stimulus interval (ISI). Standard tones were 523.25 Hz (corresponding to C5 on a piano), deviant tones were 20 cents higher (i.e., 529.33 Hz) and 20 cents lower (i.e., 517.24 Hz) with respect to the standard. The visual stimuli consisted of an image of a rectangle containing a light grey disc (see Figure 4), which subtended a visual angle of 3 degrees watched from a distance of 135 cm. The rectangle was static, yet the disc was positioned in three vertical positions: either above, below, or in the center of the rectangle, which was clearly marked by the crossing of two lines serving as a fixation cross. The displacement of the disc from the center was 0.5 degrees visual angle, i.e., small enough to be perceived as an excursion of the disc rather than as the sudden pop up of a new disc, but large enough to be clearly visible. The duration of the visual stimuli was 416 ms, i.e., there was no ISI. This was preferred because the perceptual salience of a flickering visual stimulus (i.e., one with a 316 ms ISI) much exceeded the perceptual salience of the excursion of the disc.

**Figure 4.**
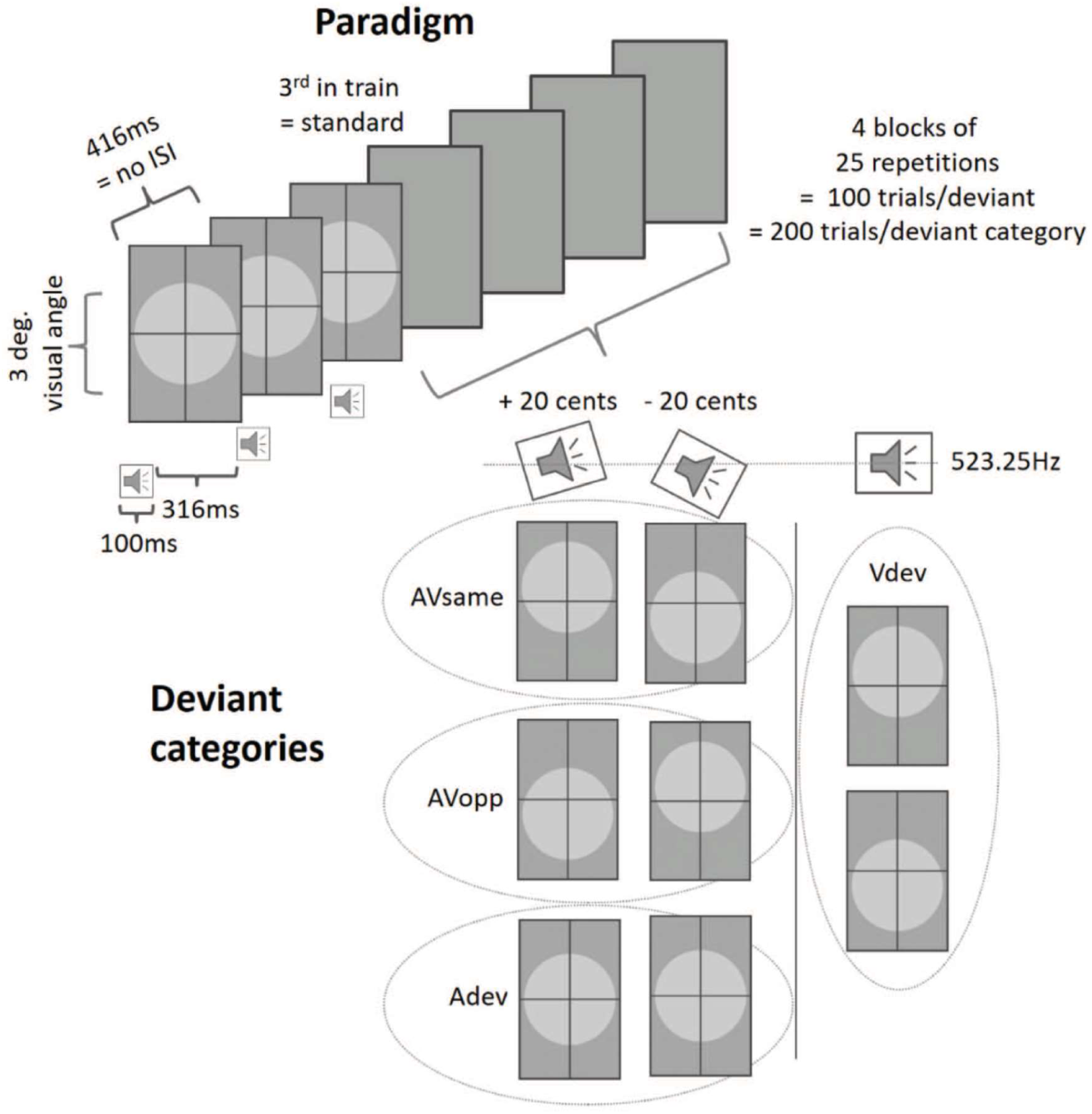
Stimuli and paradigm. The experiment consisted of a semi-attended oddball paradigm with 80% standards and eight different deviants, categorized into four deviant categories: *same direction* deviants (AVsame), *opposite direction* deviants (AVopp), *auditory-only* deviants (Adev), and *visual-only* deviants (Vdev). 12.1% of the standards (always the first in a train) were colored control task stimuli. In order for participants to stay focused on the visual stimuli, they were instructed to count one of four colors per block and report the count orally to the experimenter after each block.

The audiovisual (AV) stimuli comprised all possible combinations of the three auditory and three visual stimulus types, resulting in nine different stimulus pairs. An AV standard consisted of a standard tone coupled with a disc in center position. With higher pitch corresponding to higher vertical position, the eight AV deviants were categorized into *same direction* deviants (AVsame) (i.e., high pitch with disc above the center, or low pitch with disc below the center), *opposite direction* deviants (AVopp) (i.e., high pitch with disc below the center, or low pitch with disc above the center), and *auditory-only* deviants (Adev) (high pitch or low pitch with no excursion of the disc from the center). A *visual-only* deviant (Vdev) (no pitch change with disc below or above the center) also appeared in the paradigm. However, because of the focus on putative visual modulation of the auditory MMNm, the present paradigm was not designed to assess visual MMNm. Indeed, despite the visual only deviant included, it is not possible to extract proper vMMNm from it that can be likened to the vMMNm reported in the literature (Stefanics, Astikainen, & Czigler, 2015; Winkler & Czigler, 2012). This is the case because no actual visual standard *events* are present in a visual stream with no ISI, except on the standard immediately following the deviants, which in turn is contaminated by the response to the preceding deviant. The purpose of including the visual deviant nonetheless was to reduce predictability by ensuring that a change in the visual stream was not always accompanied by an auditory deviant.

Presentation software (Neurobehavioral systems Inc., Albany, CA, USA) was used to present the stimuli. The timing of the onset of auditory and visual stimuli was adjusted to mimic sound source perception from the perspective of the participant. This included ensuring that the sounds arrived at participants’ ears with the same delay (4 ms) as if they had been emitted from the position of the visual images themselves. The AV stimuli were presented with a stimulus onset asynchrony (SOA) of 416 ms in an oddball paradigm with 20% deviants (2.5% per deviant, 5% per deviant category). 12.1% of the remaining stimuli were control task stimuli. They were similar to the standard stimuli with respect to auditory frequency and position of the disc, yet the disc was colored: green, red, blue, yellow, all matched with respect to the luminance of the grey discs. These control task stimuli were always presented as the first stimulus after a deviant. Each of the eight deviants was presented 100 times, yet pairs of deviants that fell into the same category (Adev, AVsame, AVopp, Vdev, respectively) were concatenated in the analysis to increase the signal-to-noise ratio, yielding 200 trials per deviant category. The deviants were pseudo-randomly presented with a minimum of three and a maximum of seven standards between two deviants. As such, the distribution of the number of standards was centered on four, however, a right skew was enforced in the distribution to give relatively few instances of six and seven standards. This approached was preferred in order to reduce the predictability arising from equiprobable stimulus presentation, which may attenuate the MMNm amplitude. The total number of audiovisual stimuli was thus 4000, 800 of which were deviants. They were presented in four identical blocks preceded by 5 instances of AV standards. The duration of each block was 7 min. In addition to the four AV blocks, the experiment included two unimodal control blocks, presented as the third and fifth of the six blocks; the presentation order was counterbalanced across participants within each group. The unimodal visual control block was identical to the AV block except that the sound was turned off. In the unimodal auditory control block, which also contained 200 deviant trials (100 high, 100 low pitch trials), the visual stimuli were replaced by a silent Charlie Chaplin movie.

### Procedure

During MEG recordings, participants were seated in a magnetically shielded and sound-attenuated room with ambient lighting. Visual stimuli were projected onto a screen with a refresh rate of 60 Hz. The screen was positioned 135 cm in front of the participant. Auditory stimuli were presented via in-ear headphones (Etymotic ER•30) at a comfortable hearing level of approximately 70 dB sound pressure level (SPL). Before the experiment, a short experimental procedure varying interaural-level-difference (ILD) was administered to ensure that participants perceived the sounds as originating straight in front of them (i.e., SPL was perceived as equal in the two ears). Participants were informed about the presence of sounds and images in the experiment, and instructed to focus on the cross at the center of the screen, pay attention to the color changes of the disc and silently count the number of occurrences. There were 28 green, 32 red, 37 blue and 24 yellow discs in each block. Participants counted each color separately, and reported the count orally after each block. This task applied to the four AV blocks as well as to the visual only block in which the targets were green discs. In the auditory only block, participants were encouraged to take a break from the task and enjoy the movie. In all encounters with the participants, we referred to the auditory stimuli as “sounds” rather than “notes” or “tones” in order not to induce a semantic confound into the experiment by using a vocabulary that is often associated with the relative terms “high” and “low”. This was particularly important as all instructions were in Danish and the Danish word for “pitch” literally translates into “tone height” (*tonehøjde*). The duration of the experiment was approximately 50 minutes, including breaks between the blocks.

### Pitch threshold estimation

In a separate session, two to four weeks after the MEG session, individual pitch discrimination thresholds (PDTs) were estimated using a *two-down, one-up* adaptive staircase procedure that converges on the 70.7% performance level on the psychometric function (Levitt 1971). Using an AXB forced-choice task, the reference (X) was always a 523.25 Hz sinusoidal tone, and the task was to state whether the first (A) or the last (B) tone differed from the two other tones. The staircase terminated after 14 reversals and the threshold was calculated on the basis of an average of the last 6 reversals. Depending on participants’ responses, the duration was approximately three to five minutes. The staircase procedure was similar to the one used by Williamson, Liu, Peryer, Grierson, and Stewart (2012), yet adapted to match the stimuli and participants of the present study. The pitch threshold estimation data were also included in the analysis of a behavioral experiment reported in Møller et al., (2018).

### Musical abilities assessment

To assess the musical abilities of the participants, we used the melodic part of the Musical Ear Test (MET) (Wallentin et al., 2010). In the melodic subtest, participant were asked to judge whether pairs of melodic piano phrases were identical or not. MET scores have been shown to correlate with results of musical imitation tests typically used in auditions for music conservatories.

### MEG recording

Continuous MEG data were sampled at 1000 Hz using Elekta Neuromag Triux™ whole-head 306-channel MEG system, with 204 planar gradiometers and 102 magnetometers. This system is part of the MINDLab Core Experimental Facility at Aarhus University Hospital, Denmark. Participants’ head positions were recorded using four continuous Head Position Indicator (cHPI) coils, and bipolar surface electrodes positioned above and below the right eye (vertical EOG), at the outer canthi of both eyes (horizontal EOG), and on the right clavicle and left rib (ECG) recorded eye blinks and cardiac activity, respectively.

### MEG preprocessing

Initial preprocessing was performed using the temporal extension of the signal space separation (tSSS) technique (Taulu, Kajola, & Simola, 2004; Taulu & Simola, 2006)) implemented in Elekta’s MaxFilter software (Version 2.2.15). This involved removal of noise from electromagnetic sources outside the head and compensation for within- and between-block differences in head position as well as continuous head movement compensation using cHPI. EOG and ECG artifacts were detected and removed from the maxfiltered data with independent component analysis (ICA) using the *find_bads_eog* and *find_bads_ecg* algorithms in MNE Python (Gramfort et al., 2013, 2014). The topographies and averaged epochs of all identified EOG- and ECG-related components were visually inspected for all participants to ensure the validity of rejected components. The remaining data were band-pass filtered, using an infinite impulse response (IIR) filter with cutoff frequencies of 1 and 40 Hz.

The subsequent preprocessing was conducted using FieldTrip (Oostenveld, Fries, Maris, & Schoffelen, 2011). The MEG data were downsampled to 250 hz and separated in epochs of 516 ms starting 100 ms before the onset of the trial and ending at the onset of the following trial. Baseline correction was applied based on the 100 ms pre-stimulus window. Trials containing SQUID jumps were discarded using automatic artefact rejection with a z-value cutoff of 30.

Planar gradiometers pick up the largest signal just above a local source where the gradient has its maximum (Levänen et al., 1996; Luck, 2005). In contrast, magnetometers are more likely to also pick up activity from deeper sources and elsewhere in the cortex (Vrba & Robinson, 2002). Hence, because our focused interest was in the modulation of the neural signals emanating specifically from the auditory cortex, the sensor space analysis focused on combined planar gradiometer recordings from immediately above auditory cortex. Planar gradiometer recordings were calculated as vector sums by taking the root-mean-square (RMS) of the two gradients at each gradiometer pair resulting in a single positive RMS value for each pair.

Channel selection was accomplished through the use of the MMNm response obtained in the unimodal auditory control block. As such, the well-described MMNm response obtained in a classic oddball paradigm as used in our control block served as a functional localizer at the sensor level (Luck & Gaspelin, 2017). Initially, group-averaged difference waveforms for each channel were obtained by first subtracting each participant’s average response to the third standard after a deviant from their average response to the deviants, and then averaging across participants. Peak amplitudes and latencies of the P1m, i.e., the early, sensory-driven component which was prominent for all sounds in the present data, were identified in the standard waveforms within a 40-80 ms time window from stimulus onset. On the difference waveforms, local MMNm peaks were identified within a 100-200 ms time window following the timepoint of the P1m peak as it appeared in the standard waveform. This was accomplished using Matlab’s *findpeaks* function, which identified data samples with a greater voltage than each of its neighboring data samples. A polarity switch in the magnetometer topographies between the P1m and the MMNm components was confirmed through visual inspection of the group-averaged topographies (see Figure 5).

**Figure 5.**
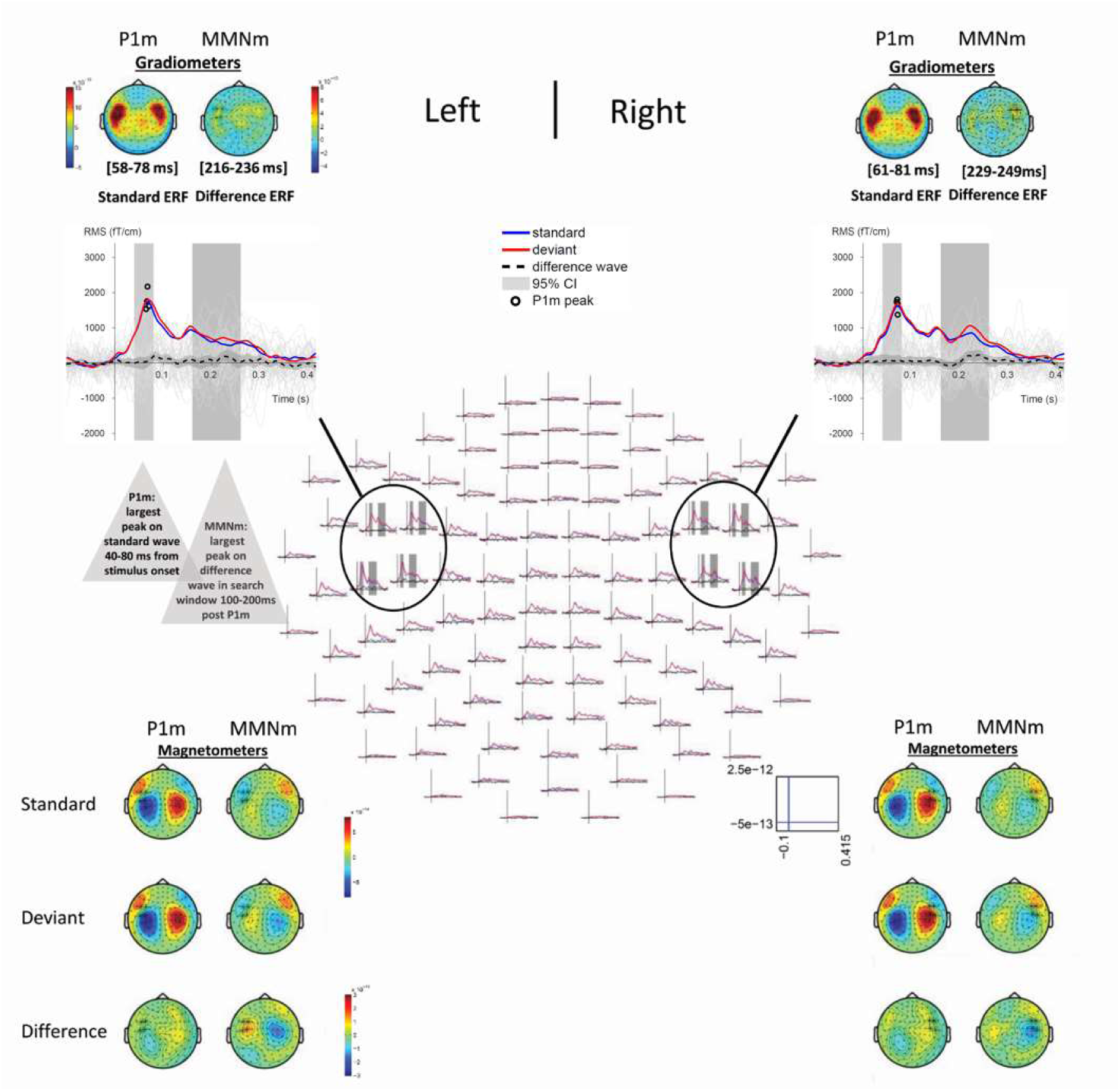
Top: Channel selection was accomplished using combined gradiometer data recorded in the unisensory auditory control block. The MMNm response obtained in this classic oddball paradigm served as a functional localizer in the spatial domain. Middle: The four channels of each hemisphere that picked up the largest group-averaged MMNm response defined the sensors-of-interest in the subsequent analyses of data obtained in the audiovisual experimental blocks. Circles in the group-averaged ERF waveforms show amplitudes and latencies of the P1m peaks in the four channels. Bottom: A polarity switch in the magnetometer data confirmed that the magnetic field of the identified MMNm peak was indeed of opposite polarity to the P1m.

MMNm amplitudes were defined as the average within a 20 ms time window centered at the MMNm peak latencies. Having identified the MMNm amplitudes for each channel in the unimodal auditory control block, all subsequent analyses were constrained to the four sensors of each hemisphere picking up the largest group-averaged MMNm response (see Figure 5) in the unimodal auditory control block. Rather than using just the peak channel, the average of four channels was used to account for individual variations in head position, size, etc. The four selected channels were also the channels with the largest P1m amplitude in the group-average standard waveform. To confirm the relevance of these four channels at the level of the individual participants, they were compared to the P1m peak channel of each participant for each hemisphere. Individual P1m amplitudes and latencies were identified within the 40-80 ms time window from stimulus onset and the P1m peak channel was defined as the channel containing the largest peak. Except for two participants whose P1m peak channel in the left hemisphere was a more posterior neighboring channel, the P1m peak channel of all participants was identical to one of the four selected channels of each hemisphere.

Due to the large inter-individual variability of the ERF difference waveforms in the experimental conditions, individual local MMNm peaks and latencies of each hemisphere and each condition were identified for each participant individually using the same procedure as described for the functional localizer, i.e., within a 100-200 ms time window following the individual participant’s P1m peak in the auditory-only block. A summary value of the MMNm amplitude was extracted for each participant, each hemisphere and each condition (see individual participants’ responses in supplementary materials, Figure S1). In cases where the MMNm peak was exceeded in amplitude by a peak which did not show opposite polarity to the P1m on the magnetometer topography, the window was narrowed further to avoid the spurious peak being picked up by the fully automated procedure. Similarly, plots of waveforms and topographies were compared across conditions within each hemisphere for each participant. Inconsistencies were typically found when MMNms were less prominent, and the search criteria were adapted when applicable to ensure consistency between the automatically selected peaks. Altogether, further constraints were necessary in one or both hemispheres for 10 participants in the unimodal auditory control block and in 1-5 out of 8 instances (4 conditions x 2 hemispheres) for 11 participants in the audiovisual experimental blocks, accounting for 9.3 % of the total number of peaks.

All statistical analyses were performed in IBM SPSS Statistics 24. Bootstrapping with 5000 samples was used to estimate 95% bias-corrected and accelerated (BCa) confidence intervals (CIs), when this more robust method was required due to skewness in the distribution of one or more variable outcomes.

## Acknowledgements

We wish to thank Signe Hagner for assistance in data collection, and Christopher Bailey for valuable technical and scientific advice at various stages of MEG recording and analyses.

