## Supplementary materials for "Poorer auditory sensitivity is related to stronger visual enhancement of the human auditory mismatch negativity (MMNm)"

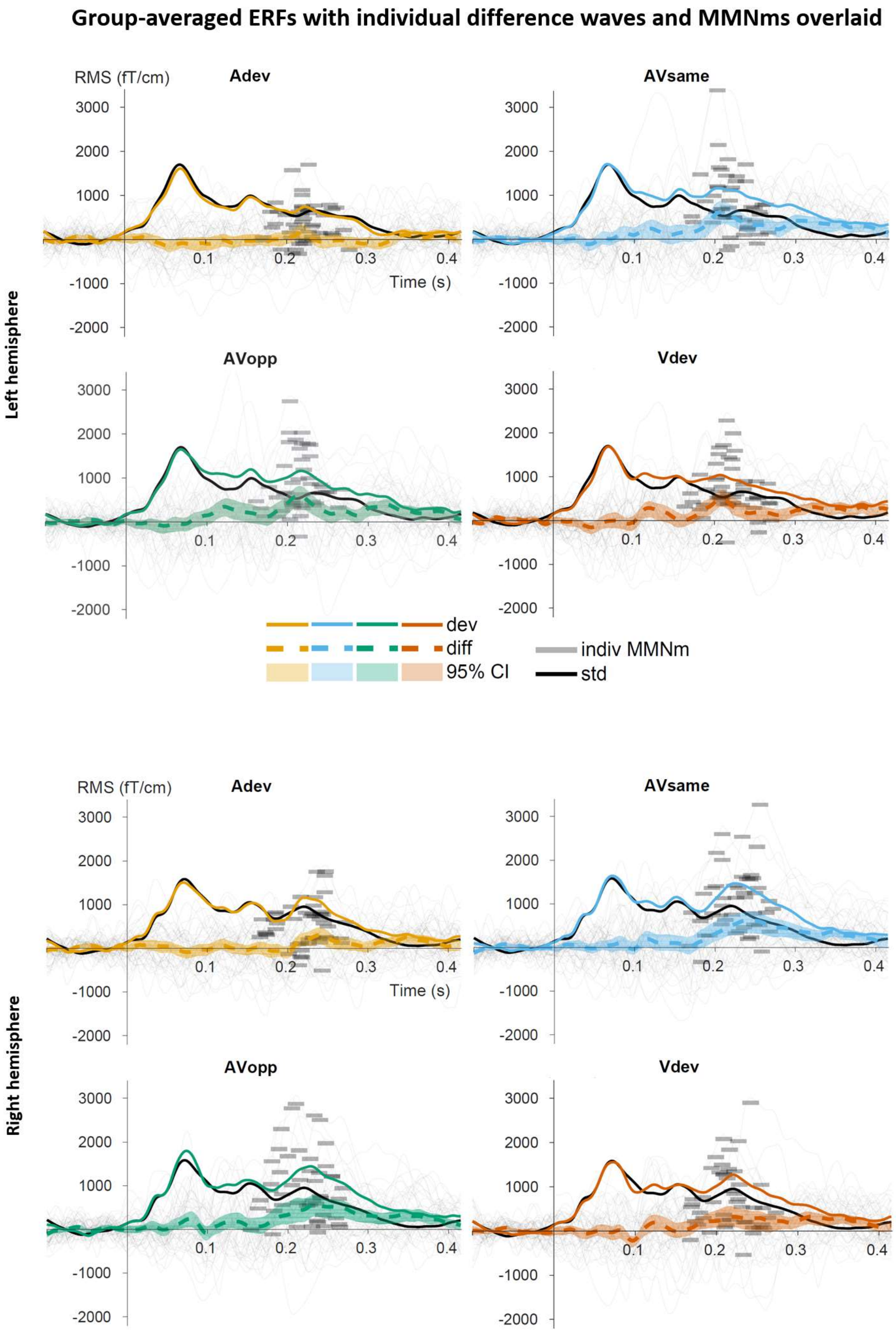


*Figure S1.* Plots show group-averaged standard, deviant and difference waveforms (incl. 95% confidence intervals) of all four conditions and both hemispheres (see corresponding topographies in Figure S2, S3, S4, and S5). Thin grey lines represent all 45 individual participants' difference waveforms. Thick grey bars show their MMNm latencies and amplitudes. The MMNm amplitudes fed into the statistical analyses were defined as a 20 ms windowed mean centered on the largest peak in a 100-200 ms window following the individual P1m latency of the standard wave obtained in the auditory-only control block.

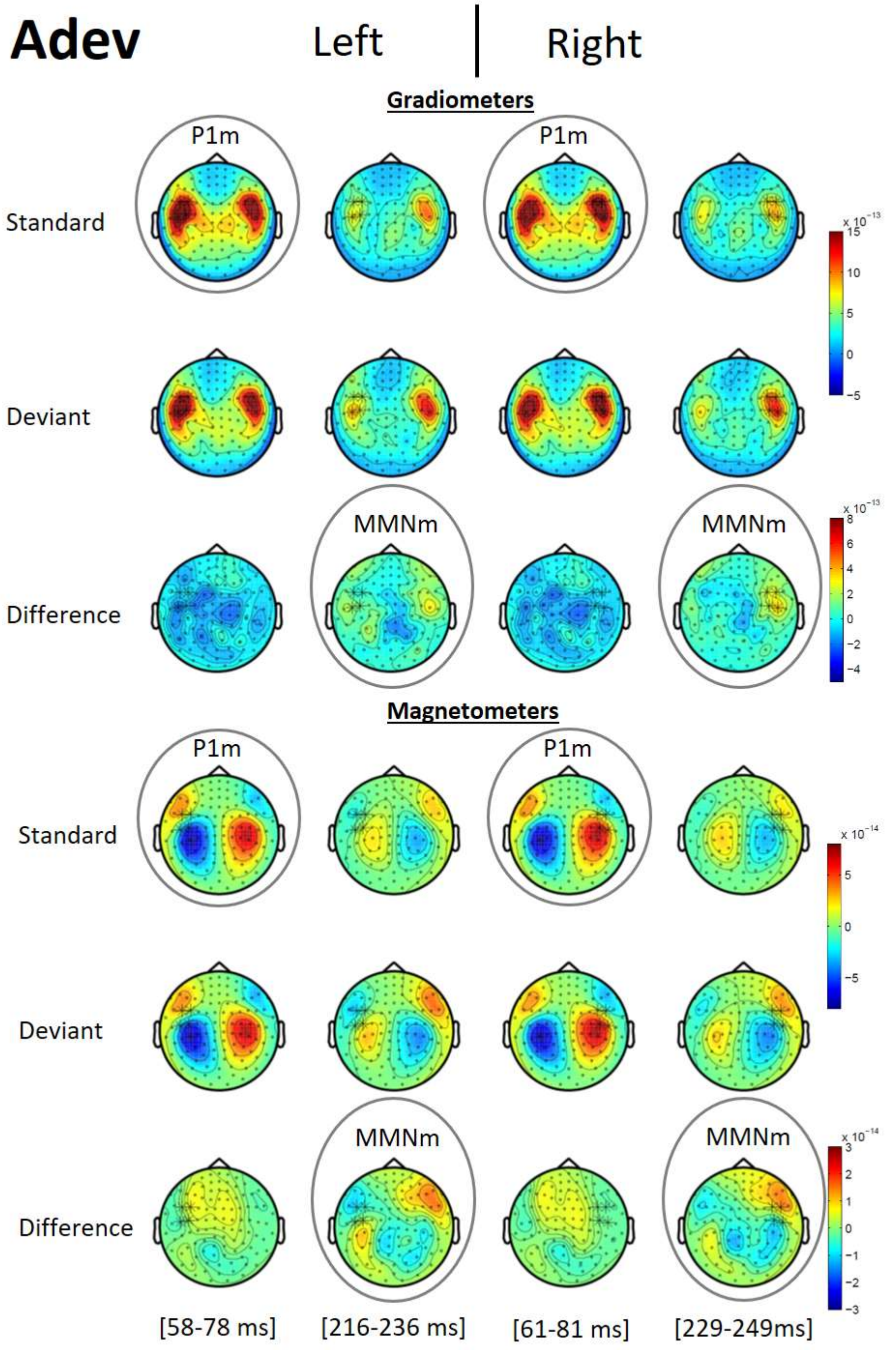


*Figure S2.* Topographic distributions of the event-related magnetic fields associated with *auditory-only* deviants (Adev), (high pitch or low pitch + no excursion of the disc from the center). Topographies show 20 ms windows centered on the peak latency of the grand mean peak of all four conditions. Note, that statistics were performed on the amplitudes of MMNm responses identified in each individual participant (see individual responses to each deviant in Figure S1).

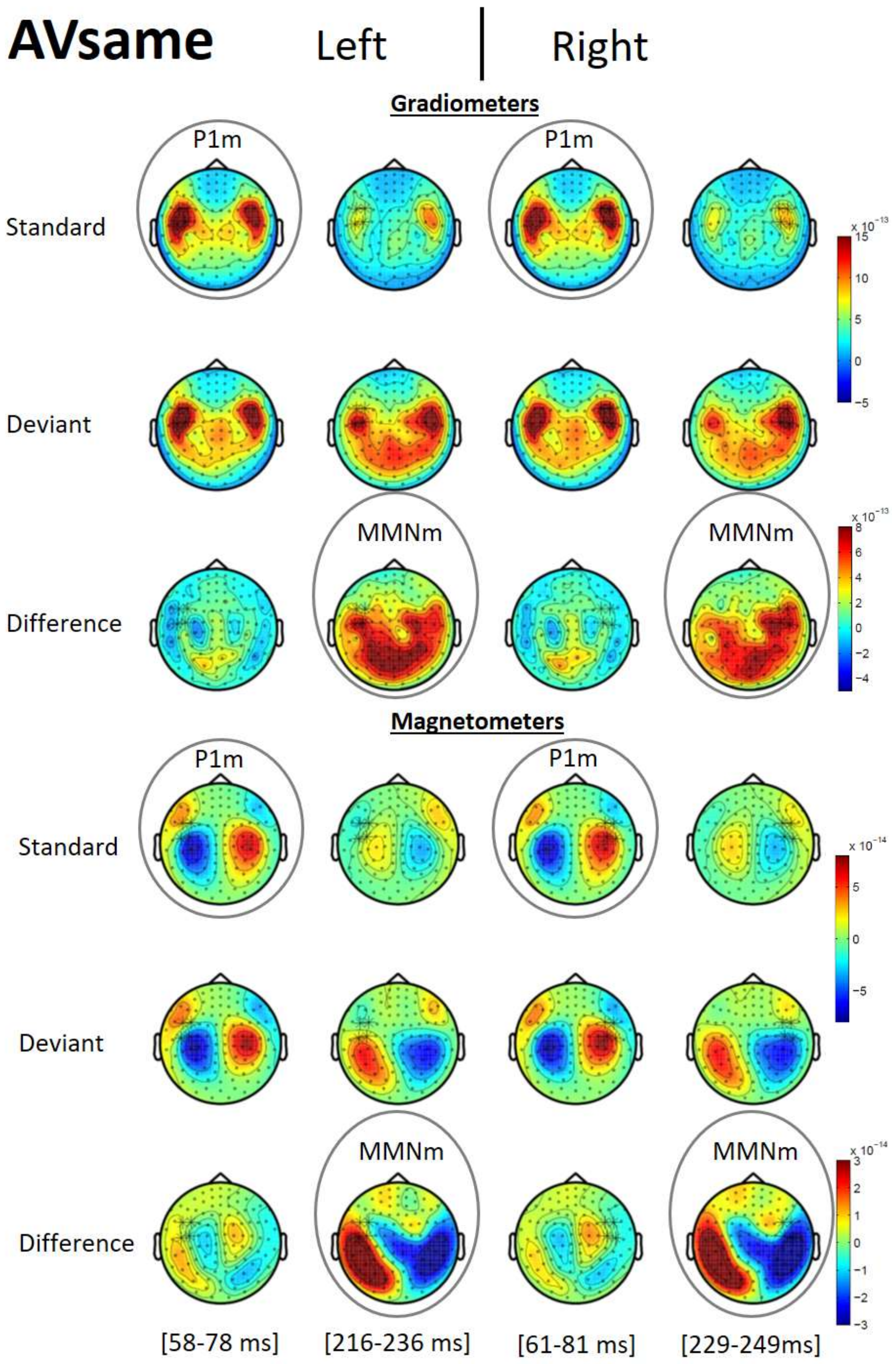


*Figure S3.* Topographic distributions of the event-related magnetic fields associated with *same direction* deviants (AVsame), (i.e., high pitch + disc above the center, or low pitch + disc below the center). Topographies show 20 ms windows centered on the peak latency of the grand mean peak of all four conditions. Note, that statistics were performed on the amplitudes of MMNm responses identified in each individual participant (see individual responses to each deviant in Figure S1).

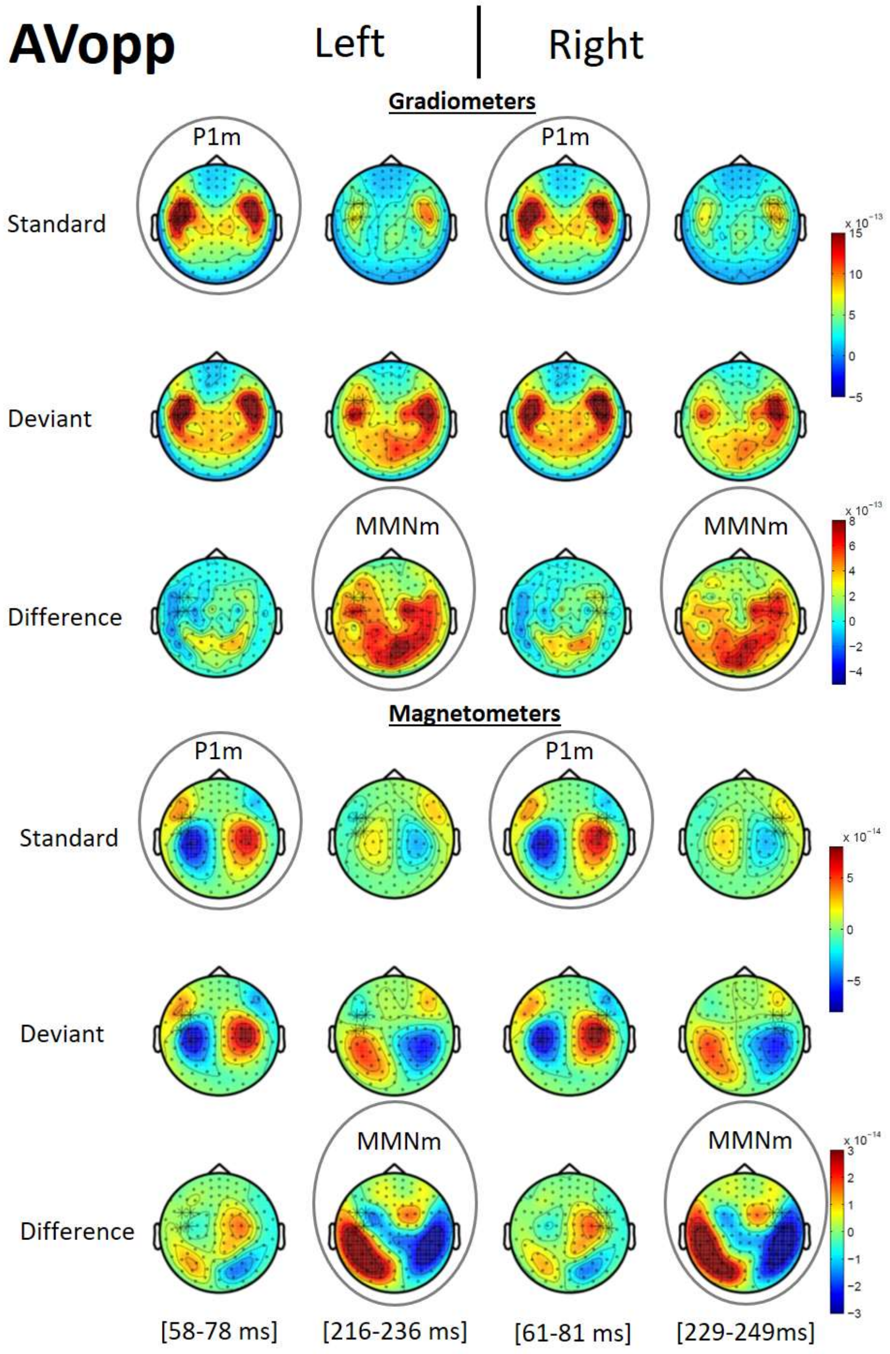


*Figure S4*. Topographic distributions of the event-related magnetic fields associated with *opposite direction* deviants (AVopp), (i.e., high pitch + disc below the center, or low pitch + disc above the center). Topographies show 20 ms windows centered on the peak latency of the grand mean of all four conditions. Note, that statistics were performed on the amplitudes of MMNm responses identified in each individual participant (see individual responses to each deviant in Figure S1).

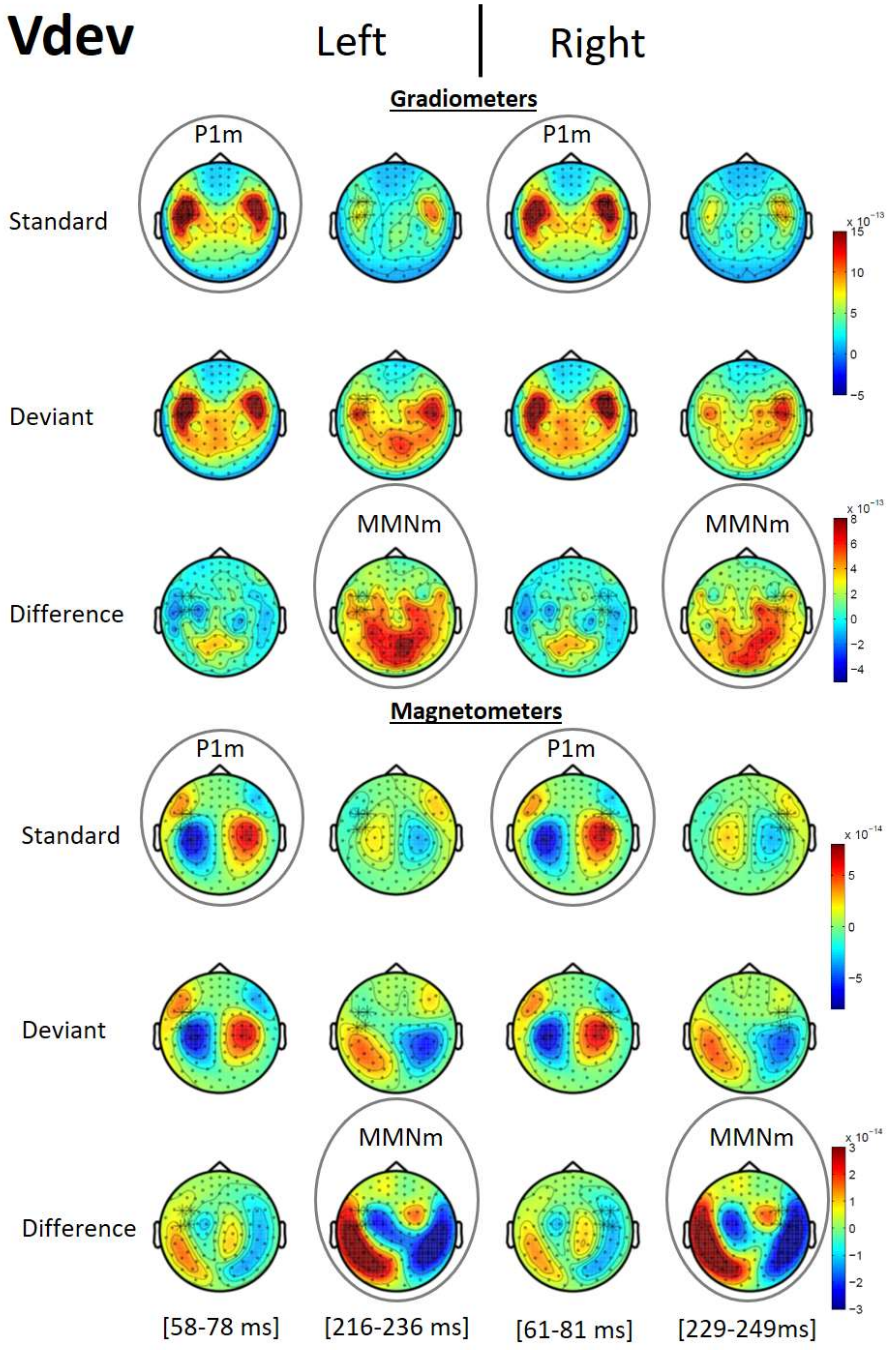


*Figure S5*. Topographic distributions of the event-related magnetic fields associated with *visual-only* deviants (Vdev), (no pitch change + disc below or above the center). Topographies show 20 ms windows centered on the peak latency of the grand mean of all four conditions. Note, that statistics were performed on the amplitudes of MMNm responses identified in each individual participant (see individual responses to each deviant in Figure S1).

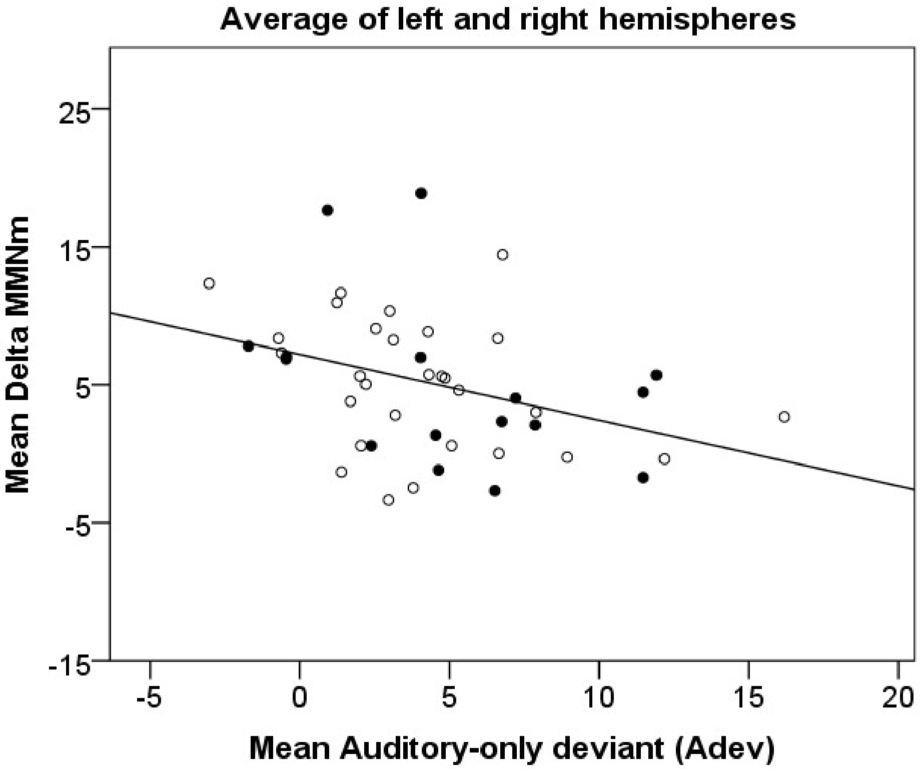


*Figure S6.* The analyses reported in Figure 4 of the manuscript were performed separately in the two hemispheres. This scatterplot depicts the averaged responses across hemispheres within participants. The plot shows participants' average visually induced enhancement in MMNm amplitude, mean ΔMMNm, as a function of their average MMNm amplitude elicited in response to auditory deviants, i.e. without complementary visual cues. The correlation is significant as assessed with bootstrapped bias-corrected and accelerated (BCa) 95% confidence intervals (5000 samples), $r$ = -.367, 95% CI [-.564, -.167].
